# The Privileged Link Between Number and Space: Day-old Chicks (*Gallus gallus*) Show Spatial-Numerical but Not Spatial-Quantity Associations

**DOI:** 10.64898/2026.08.03.742501

**Authors:** Arianna Felisatti, Lucia Regolin, Rosa Rugani

**Author notes:** Correspondence concerning this article should be addressed to Arianna Felisatti and Rosa Rugani. Data are publicly available at: https://osf.io/5eu4x/overview?view_only=316d033fb4c54564b9d6e0dc6d120b95.

## Abstract

Spatial-numerical association reflects an internal “mental number line” where numerosities are mapped from left to right in space. While evidence from 8- to 9-month-old human infants suggests a privileged link between number and space, these findings do not definitively establish whether discrete numerosity or continuous quantity is the primary driver of spatialization due to the relatively extensive postnatal experience. Using domestic chicks (*Gallus gallus*), a model testable with minimal postnatal experience, we investigated precocial predispositions to map both discrete numerosities and continuous quantity (physical size) onto space. In Experiment 1, we replicated spatial-*numerical* association: chicks associated relatively smaller numerosities with the left hemispace and larger ones with the right hemispace. In Experiment 2, however, spatial-*quantity* association for physical size was asymmetric: chicks showed a rightward preference for smaller sizes and no congruent small-left/large-right mapping. Together with infant data, these findings indicate that a generalized and congruent spatial-magnitude association does not emerge at the earliest stages of development, and challenge accounts positing that spatial mappings for continuous magnitudes precede and underpin the spatial organization of number.

## Introduction

For centuries, mathematics was regarded as a uniquely human *logos*, inextricably tied to language and formal education. Modern comparative research has questioned this anthropocentric boundary, revealing a continuum between human and animal cognition in numerical abilities. Numerical information is embedded in the natural world such as in amount of food sources, number of group members or predators (Piantadosi & Cantlon, 2017). Successful organisms must constantly process numerical information to compete and survive (Dudine et al., *Under Review*; Nieder, 2026). Animals are endowed with a number sense that supports efficient decision-making across ecologically relevant contexts (Giurfa, 2019; Macchinizzi et al., 2025; Regolin et al., 2025; Reznikova & Ryabko, 2011; Rugani, 2018). All these survival-related scenarios unfold within a physical environment where items and agents are naturally distributed in space. Given that these quantities are encountered through spatial interaction, the brain must integrate numerical magnitude with spatial coordinates to guide effective action.

Notably, evidence shows that numerical processing is tightly linked with space. Humans and animals typically associate small numerosities with the left and large numerosities with the right (Dehaene et al., 1993; Galton, 1881; Restle, 1970; reviews in Felisatti et al., 2026; Toomarian & Hubbard, 2018). Spatial-numerical association (SNA) has been documented across a wide range of species and developmental stages: in human adults (Nemeh et al., 2018; Porru et al., 2025; Rugani et al., 2017; Rugani & Sartori, 2016; Zhou et al., 2016), children (De Hevia & Spelke, 2009; Rugani, Zhang, et al., 2022; Straulino et al., 2026), preverbal newborns (Di Giorgio et al., 2019; Volpi & De Hevia, 2026), primates (Adachi, 2014; Annicchiarico et al., 2026; Drucker & Brannon, 2014; Gazes et al., 2017; Johnson-Ulrich & Vonk, 2018; Rugani et al., 2024; Rugani, Platt, et al., 2022; but see Beran et al., 2019), birds (Felisatti, Macchinizzi, et al., 2026; Rugani et al., 2010, 2015a, 2020, 2025), fish (Potrich et al., 2026; but see Triki & Bshary, 2018), and insects (Giurfa et al., 2022). SNA has been interpreted as evidence of an internal “mental number line” defined as an analogue, horizontally oriented continuum where numerical magnitudes are mapped from left to right. Increasing comparative studies supports the consideration of SNA as an ancient, conserved feature of animal cognition that precedes the influence of learning-related aspects and contingent situations (Eccher et al., 2026; Felisatti, Giurfa, et al., 2026).

In the physical world, numerosity is intrinsically correlated with multiple continuous variables, including cumulative surface area, total perimeter, volume and density, as described by A Theory of Magnitude (ATOM), which proposes that space, time, and quantity share a generalized magnitude-processing system grounded in the parietal cortex (Bueti & Walsh, 2009; Walsh, 2003). While many studies have controlled for these continuous variables and found SNA independent of them (Di Giorgio et al., 2019; Giurfa et al., 2022; Rugani et al., 2015a), other research has noted that SNA may only emerge when animals can utilize non-numerical cues (Annicchiarico et al., 2026; Johnson-Ulrich & Vonk, 2018; Potrich et al., 2026). Indeed, spatial associations are not restricted to discrete numerical magnitude but extend to various

*prothetic* dimensions, namely those defined by “more than” or “less than” variations, such as volume (Kirjakovski & Utsuki, 2012), weight (Dalmaso & Vicovaro, 2019), speed (Vicovaro et al., 2025), temporal duration (Di Bono et al., 2012), face age (Dalmaso et al., 2026) and luminance (Loconsole et al., 2021). A recent meta-analysis on human studies confirmed spatial associations for continuous quantities (known as Spatial-Quantity Association of Response Codes, SQUARC), although weaker than those for discrete numerosities (known as Spatial-Numerical Association of Response Codes, SNARC) (Macnamara et al., 2018).

However, because numerosity naturally covaries with continuous magnitudes, it has been argued that spatial–numerical association may not be specific to discrete numerosity but instead reflect a more general mapping between space and quantities. Specifically, Leibovich and colleagues (Leibovich et al., 2017) proposed a shift from a “number sense” to a “magnitude sense” view, according to which discrete numerosity is not an innate primitive but is instead derived through development by learning the natural correlations between number and continuous magnitudes in the environment. A clear prediction follows from this model: if spatial mappings are grounded in a domain-general representation of magnitude, then continuous dimensions such as physical size should exhibit a congruent small-left/large-right mapping, mirroring the spatialization of discrete numerosities.

Initial insight into the developmental trajectory of these associations comes from studies on human adults and infants (Bulf et al., 2014, 2016). Specifically, they employed Posner-like cueing tasks to measure whether numerosities and physical size orient visual attention across the horizontal space. Human adults demonstrated that both small numerosities and physical size facilitated the detection of targets appearing on the left side while larger numerosities and physical size facilitated the detection of targets on the right side of the screen (Bulf et al., 2014). Conversely, 8-to 9-month-old infants reported significant lateralized attentional shifts only with numerosities and not with physical size (Bulf et al., 2016). Surprisingly, physical size led to a non-significant opposite trend, with longer time to look at the left-side target after a small-size image and at the right-side target after a large-size image. The authors interpreted these findings as indicative of a “privileged link” between number and space that only later generalize to physical size through ontogenetic experience. However, since infants at this stage have already accumulated over eight months of postnatal experience, it is difficult to definitively disentangle innate biological predispositions from associations refined or acquired through environmental interaction and scanning routines. Consequently, testing day-old populations with minimal postnatal experience is essential first to determine whether the mapping of continuous quantity onto space is functional from birth, and secondly to determine its evolutionary functionality.

The present study addresses this gap by exploring whether day-old domestic chicks exhibit spontaneous spatial association for physical size that aligns with their documented numerical mapping. Domestic chicks are as an ideal model species as they can be tested a few days after hatching with minimal postnatal experience. If spatial mappings reflect a domain-general representation of magnitude, chicks should associate smaller sizes with the left side and larger sizes with the right, paralleling their documented mapping of smaller and larger numerosities onto left and right space, respectively.

## Materials and Methods

All experimental procedures were approved and conducted in strict adherence to the guidelines provided by the Committee for Animal Welfare of the University of Padua and the Ministry of Health of the Italian Republic (Prot. N. 729/2016-PR, 22/07/2016 and Prot. N. 306/2019-PR, 14/04/2019). This comprehensive compliance addressed both national and European directives concerning animal research.

### Subjects

A priori power analysis was conducted using G*Power 3.1 to determine the required sample size for a one-tail Wilcoxon signed-rank test (matched pairs). Based on Cohen’s guidelines (Cohen, 1988), a medium effect size was anticipated (*d*_*z*_ = 0.50). With an alpha level of .05 and a statistical power of 80%, the analysis indicated that a sample size of 24 chicks was required. This is in line with the sample size of previous studies assessing SNA in day-old chicks (Rugani et al., 2015a, 2020).

A total of forty-six male domestic chicks (*Gallus gallus domesticus*) were trained and tested. They were obtained as fertilized eggs from local hatcheries (Agricola Berica, Montegalda, Vicenza, Italy, or Società Agricola La Pellegrina Spa, San Pietro in Gù, Padova, Italy) and hatched in the laboratory. Twenty-four chicks participated in Experiment 1 (testing spatial association of discrete numerosity), while twenty-two chicks took part in Experiment 2 (testing spatial association of continuous quantity). Post-hatching, chicks were housed in social groups of two or three within open-top metal cages. Environmental parameters were strictly controlled, with a temperature of 28–31 °C and humidity of 68%. In the rearing room, fluorescent lamps were positioned 15 cm above each cage. Birds were provided with chicks’ crumble and water *ad libitum*. In addition, twice a day they were offered a few mealworms (*Tenebrio molitor* larvae) which served as food reward during training.

Experimental procedures started on Day 3. Food was removed from the cage to enhance motivation and, two hours later, chicks underwent a *shaping* phase functional to make them learn to circumnavigate a central panel within the apparatus. After shaping and a two-hour rest in their rearing cages, chicks completed their first training session which was immediately followed by Test 1. After an additional one-hour rest, subjects underwent a second training session and Test 2. At the end of the experimental activities, subjects were placed back into social groups and donated to local registered farmers. The experimental timeline is outlined in **Table 1**.

**Table 1.** Outline of experimental procedures.

| Time | Procedures |
| --- | --- |
| **Day 1, morning** | Housing in standard conditions<br>(within 24 hours from hatching) |
| **Day 2, all day** | Standard rearing conditions |
| **Day 3, from 8 a.m. to 10 a.m.** | Removal of food jars (2 hr before shaping) |
| **Day 3, from 10 a.m.** | Shaping + 1 hr rest + Training session 1 + 1 hr rest |
| **Day 3, early afternoon** | Test 1 + 1 hr rest |
| **Day 3, mid afternoon** | Training session 2 + Test 2 |

### Apparatus

The same experimental apparatus was used for shaping, training and test. The setup was housed in a controlled environment with temperature of 25°C and humidity of 70%. The lights were provided with four 58 W ceiling lamps. The apparatus consisted of a diamond-shaped arena made up of white plastic panels, 20 cm high (**Figure 1**). A transparent, removable partition (10 × 20 cm) was situated approximately 10 cm from the arena’s main vertex to define the starting area. Each subject was held within this area for five seconds before the start of each trial to allow visual inspection of the stimuli and the arena before being released. During inter-trial intervals, subjects were gently moved to an adjacent opaque box (20 × 40 × 40 cm) to prevent them from seeing the experimenter changing the stimuli. The stimuli were presented on panels (16 × 8 cm) with 3-cm bent-back sides, which ensured that chicks could not see the food reward (a mealworm) hidden behind the panel before circumnavigating it. In the training phase, a single panel was placed in the center of the arena, 40 cm directly in front of the starting area. At test, two identical panels were positioned symmetrically, 30 cm apart, to the left and right of the main vertex. To facilitate the scoring of chicks’ choices for either panel, a vertical partition was located in between the panels so as to create to identical sectors.

**Figure 1.**
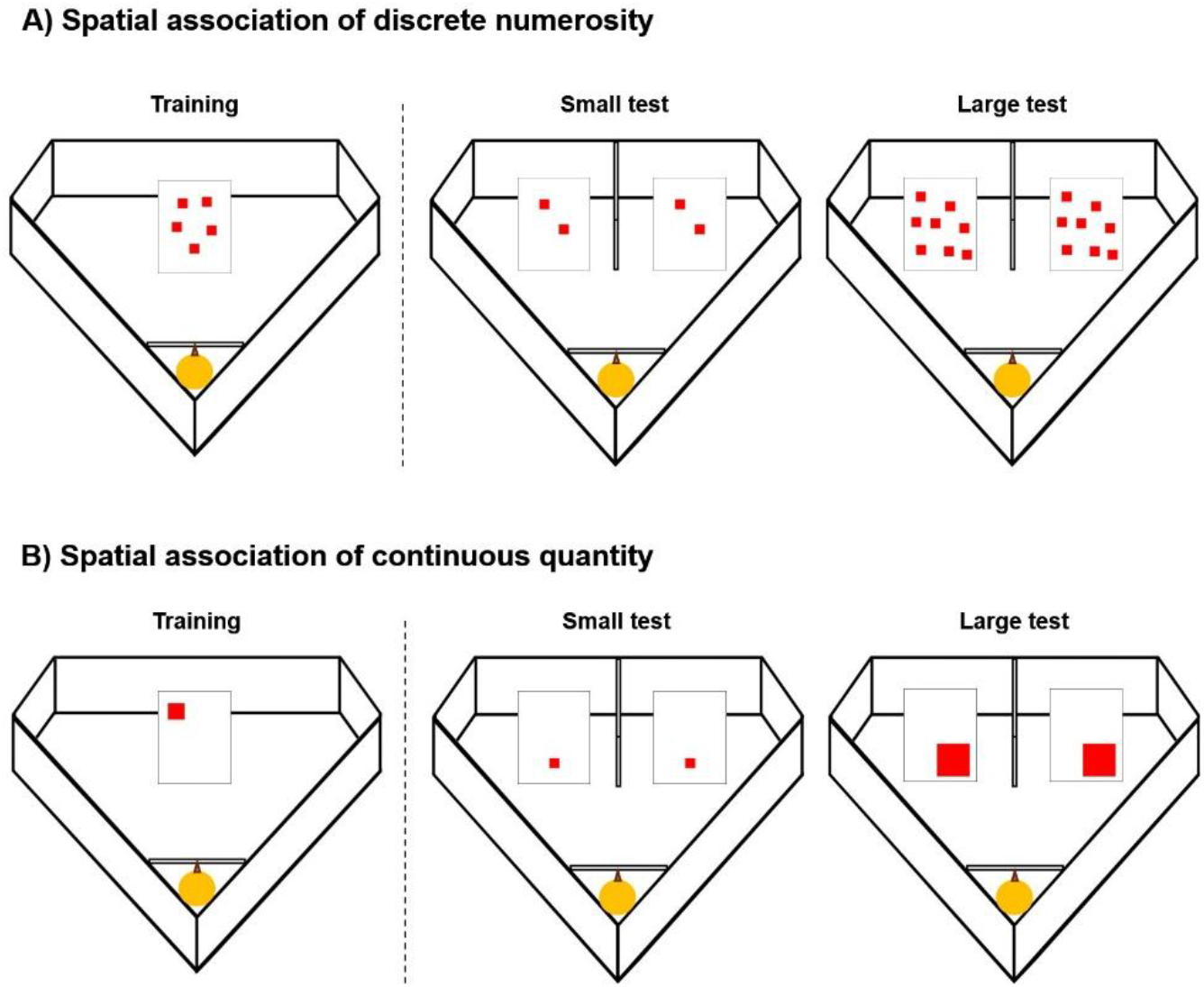
Experimental apparatus and example of the stimuli used in the training and test in Experiment 1 (A) and 2 (B).

### Stimuli

Training and test stimuli consisted of static 2D images printed on white rectangular boards (11.5 × 9 cm).

In Experiment 1, the stimuli depicted a specific number of red squares (each square measuring 1 × 1 cm). In each trial, a stimulus (during training) or a pair of stimuli (during test) were placed on the panel(s) (**Figure 1A**). The training stimuli depicted five red squares. To prevent cueing due to the spatial arrangement of the squares, 20 different training stimuli (one for each training trial) were created, featuring randomized spatial arrangements of the squares with inter-element distances ranging from 0.3 to 3.8 cm. The test stimuli depicted either 2 identical red squares (Small test) or 8 identical red squares (Large test; **Figure 1A**). Five test stimuli, differing for the spatial arrangement of the squares, were used for each test (Small test: 2 vs. 2; Large test: 8 vs. 8). Two identical copies of each test stimulus were printed and were simultaneously presented at test. All stimuli were designed following an established paradigm used in previous studies (Rugani et al., 2015a, 2020).

In Experiment 2, the stimuli depicted a single red square. The training stimuli consisted of a red square of intermediate size (2 × 2 cm). Twenty training stimuli (one for each trial) differing in the spatial arrangement of the single square on the white board were created. The test stimuli displayed a single square either smaller (1 × 1 cm; Small test) or larger (4 cm × 4 cm; Large test; **Figure 1B**) as compared to the training one. Five different test stimuli with different spatial disposition of the square were produced for each of the two tests. At test, two identical copies of a same stimulus were presented.

### Experimental phases

On the morning of Day 3, each subject underwent ***shaping***, during which it was acquainted to moving and feeding in the apparatus. In the first stage of shaping, the chick familiarized with the environment and a single panel was present. Specifically, each subject was first allowed to explore the arena and was then guided to approach the central panel by dropping a mealworm progressively farther from the starting area and closer to the panel, across five subsequent trials. In the final stage of shaping, the chick learned to search for food *behind* the panel. While confined in the starting area, the chick observed a mealworm being placed in front of the panel (long tweezers were always used to handle the mealworms) and then being moved behind it. The mealworm disappeared first to the right and then to the left of the panel, alternating directions. Once released, the bird could search for the mealworm behind the panel. Shaping was successfully concluded when the subject confidently moved from the starting area to the area behind the panel and eat the reward for three consecutive trials.

Following shaping, subjects began the formal ***training*** phase. In each trial, a single stimulus was placed on the central panel. The chick was confined in the starting area for five seconds and then released into the arena to find the food reward in a maximum of one minute. Training was considered successfully completed when the subject managed to circumnavigate the panel on 20 consecutive trials, each of which was reinforced with a mealworm. Chicks that reported little motivation or lack of interest in the food reward were excluded from the study (approximately 25% of the subjects). Only those chicks that successfully met the training criterion proceeded to the tests and were included in the final dataset.

The ***Test*** phase comprised a Small test and a Large test. Each of these consisted of five non-reinforced trials. At the beginning of every trial, the chick was held in the starting area, behind the transparent partition, for five seconds. In front of the starting area the two panels were present for the first time. Each of them displayed a copy of the same stimulus (identical number, size and spatial disposition). Once removed the transparent partition, the chick was free to walk anywhere within the arena for up to one minute. A trial was considered concluded as soon as the subject circumnavigated one of the two panels. A valid choice was defined as the moment the chick’s head and at least ¾ of its body entered the area behind a panel (Felisatti, Macchinizzi, et al., 2026; Rugani et al., 2015a, 2020). Only one choice was allowed and scored per trial. During the inter-trial interval (approximately 15 seconds), the subject was placed in an opaque box while the experimenter swapped the panels and changed the stimuli. If a chick failed to express a choice within one minute, the trial was repeated immediately for up to three times. This procedure continued until each subject completed two full sessions of five valid trials each. To prevent the experimenter’s presence from influencing behavior, once the chick was positioned within the arena and the transparent partition was removed, the experimenter stepped away from the arena and observed the trial via a monitor connected to the videocamera. Test trials were video-recorded for subsequent analysis.

## Results

For each experiment and test, the mean of choices for the left panel (%) was arbitrarily chosen for each chick following the formula: (number of left choices/total number of trials) ×100. Scores could range from 0% (left panel never chosen) to 100% (left panel always chosen).

We used non-parametric statistics: The Mann-Whitney test to analyse differences between groups, and the Wilcoxon signed-rank test to analyse differences between Small and Large tests and from chance level (50%). We reported the effect size as the rank-biserial correlation (rrb).

In addition, to rule out the possible influence of trial repetitions on the results, we ran a General Linear Model (GLM) on the choice made by the chick in the very first trial (Rugani et al., 2015b) with a binomial family and logit link function. Test (Small vs. Large) was included as a fixed factor.

All analyses were conducted using JASP 0.95.1.0 (JASP Team, 2026).

Data are publicly available at:

https://osf.io/5eu4x/overview?view_only=316d033fb4c54564b9d6e0dc6d120b95

### Experiment 1: Spatial association of discrete numerosity

#### Five trials

A Mann-Whitney test on the percentage of choices for the left panel did not reveal any difference between chicks that underwent the Small test as first (n = 12) or as second (n = 12) (U = 95.50, *p* = .18, rrb = -0.326). Data were therefore merged. A two-tail paired-samples Wilcoxon signed-rank test revealed that the Small test led to percentages significantly larger than the Large test (n = 24 chicks, W = 205.0, *p* = 0.002; rrb = 0.775; **Figure 2A**). A Wilcoxon signed-rank test was then performed to compare the data against chance level (50%). In the Small test, chicks preferred the left panel (n = 24 chicks, mean = 60.83%, SE = 25.35, V = 218.50, *p* = 0.047; rrb = 0.457), while in the Large test, they preferred the right panel (n = 24, mean = 37.50%, SE = 24.54, V = 75.00, *p* = 0.030; rrb = -0.500).

**Figure 2.**
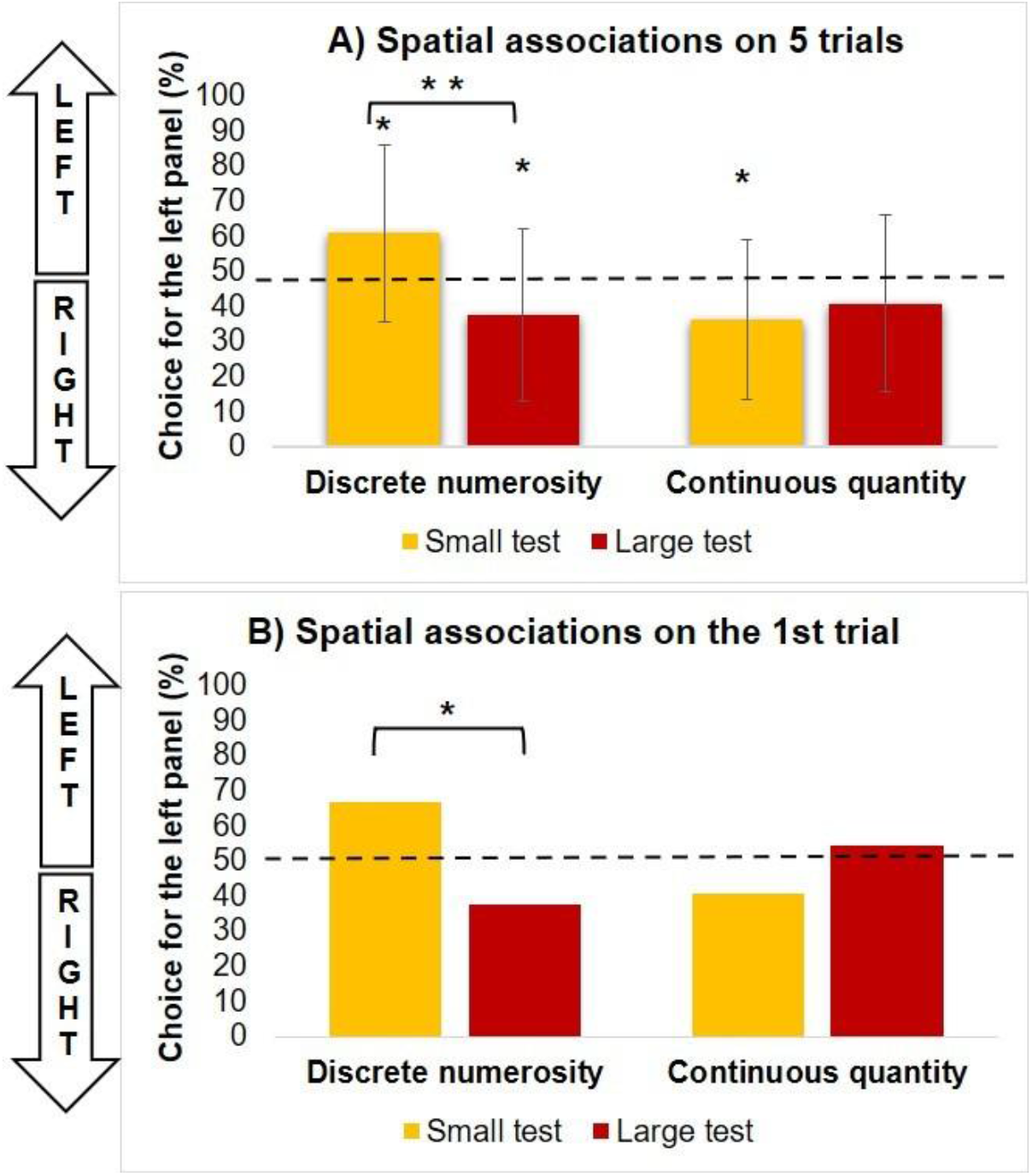
Mean of choice for the left panel (%) as a function of Test (Small test vs. Large test) when considering 5 trials (A) or only the first trial (B). The horizontal line indicates chance level (50%). Asterisks refer to significant differences from chance level within each test and differences between tests using the Wilcoxon test, (*: p < .05; **: p < .01).

#### First trial

The GLM revealed a significant effect of Test (χ^2^(1) = 4.151, *p* = .042; **Figure 2B**), confirming the results even when considering the first trial only. Small test led to a higher percentage of left choices (mean = 66.66%) compared to the Large test (mean = 37.5%).

### Experiment 2: Spatial association of continuous quantity

#### Five trials

A Mann-Whitney test on the percentage of choices for the left panel did not reveal any difference between chicks that underwent the Small test as first (n = 10) or as second (n = 12) (U = 34.50, *p* = 0.89, rrb = 0.425). Data were merged and submitted to a two-tail paired-samples Wilcoxon signed-rank test which reported absence of a significant difference between the Small and Large tests (n = 22 chicks, W = 47.0, *p* = 0.746, -0.105; **Figure 2A**). To assess differences with respect to the chance level (50%), a Wilcoxon signed-rank test was performed. In the Small test, chicks preferred the right panel (n = 22 chicks, mean = 36.36%, SE = 22.79, V = 54.00, *p* = 0.016; rrb = -0.573), while in the Large test, they did not show a significant preference for either the panel (n = 22, mean = 40.91%, SE = 25.05, V = 78.50, *p* = 0.115, rrb = -0.379).

#### First trial

Performance at the tests on the first trial did not reach significance (χ^2^(1) = 1.217, *p* = .270; **Figure 2B**), confirming the overall results. Nevertheless, in the first trial there was a trend for smaller percentages in the Small test (mean = 40.91%) compared to the Large test (mean = 54.55%).

## Discussion

Our study aimed to clarify whether spatial mappings for continuous quantity arise as early and as coherently as those for discrete numerosity. Although infant data have suggested a privileged link between number and space (Bulf et al., 2016), they cannot be conclusive about the origin of this link because infants are tested after several months of life. By testing a precocial species with minimal postnatal experience, we show that 3-day-old chicks already exhibit a robust small-left/large-right mapping for numerosities, but no corresponding congruent mapping for physical size. This dissociation indicates that the spatial organization of number emerges earlier and more consistently than spatial-quantity association, challenging strong versions of “sense of magnitude” accounts (Leibovich et al., 2017; Walsh, 2003) and supporting the view that number–space mappings may provide a biological scaffold onto which other magnitudes are integrated over development.

In two experiments, we used an established behavioural paradigm with 3-day-old domestic chicks aimed at replicating the spatial association for discrete numerosity (Experiment 1) and testing a corresponding spatial association for physical size (Experiment 2).

In Experiment 1, we successfully replicated the spatial–numerical association in young chicks, showing that day-old, naïve chicks associate smaller numerosities with the left and larger ones with the right space (Rugani et al., 2015a, 2020). When tested with a numerosity smaller than the training one (2), chicks showed a significant preference for the left panel, conversely, when tested with a larger numerosity (8), they preferred the right panel. The effect remained robust even when considering only the first trial of each test (Rugani et al., 2015b). Since previous studies had already shown that SNA in chicks persists when continuous variables are controlled (Rugani et al., 2015a), in Experiment 1 we selected stimuli in which discrete numerosity covaried with continuous quantitative dimensions. This provided a benchmark for the expected small-left/large-right mapping when numerical and quantitative information were both available and congruent, before exploring in Experiment 2 whether physical size alone could prompt a corresponding spatial mapping.

Experiment 2 provided no evidence for small-left and large-right associations: When bilaterally exposed to smaller sizes, chicks circumnavigated the right panel. This is opposite with what expected from a generalized magnitude system where small numerosities and small physical size should produce a bias in the same direction, namely to the left; whereas large numerosities and large physical size should produce a rightward bias. However, our results mirror the trend found in preverbal infants, who exhibited a significant association between small numerosities and the left side and large numerosities and the right side, together with a non-significant trend in the opposite direction for the physical size, with smaller sizes associated with the right and larger sizes with the left (Bulf et al., 2016).

How can these patterns be interpreted? Currently, three models have been advanced to explain the origin of SNA in preverbal and non-verbal population, each making distinct predictions about the presence and direction of spatial association of continuous quantity.

The <u>Emotional-Valence Model</u> (Vallortigara, 2018) postulates that smaller numerosities are intrinsically linked to negative emotions which, engaging the right hemisphere, induce leftward behavior. Vice versa, larger numerosities activate positive emotions, which engage the left hemisphere (Davidson, 2004), and guide behavior towards the right. While this model offers a potential explanation for SNA, it faces criticism regarding its ecological validity because in nature the emotional valence of numerosity is often highly dependent on context (e.g., a larger amount of food is positive, but a larger number of predators is negative). Additionally, if the same reasoning is applied to the physical size, smaller/larger images triggering negative/positive emotions, would result in leftward/rightward behavior, respectively. Instead, day-old chicks demonstrate a significant rightward preference when presented with smaller-size images.

The <u>Right Hemisphere Dominance Model</u> (Rugani et al., 2016, 2025; Rugani, Platt, et al., 2022; Rugani & Regolin, 2020) proposes that the internal organization of magnitude is rooted in the functional lateralization of the vertebrate brain, where the right hemisphere is specialized for both spatial and numerical processing. Because fibers cross at the optic chiasm, this specialization induces a contralateral (leftward) attentional shift (Deng & Rogers, 1998; Reuter-Lorenz et al., 1990; Rogers & Vallortigara, 2021). This biological bias, acting as an anchor, leads organisms to explore the environment from left to right, with consequent association of smaller numerosities (encountered first) with left and larger numerosities with right. This model well accounts for the “privileged” link between numerical magnitude and space and predicts the absence of spatial preferences for non-numerical magnitudes, such as physical-size, in light of the lack of numerical processing required with consequent involvement of the right hemisphere.

The <u>Brain’s Asymmetric Frequency Tuning Model</u> (BAFT; Felisatti, Laubrock, et al., 2020; see also Felisatti, Aagten-Murphy, et al., 2020) is inspired by hemispheric specialization for spatial frequency processing. Spatial frequencies are low-level visual features extracted from any visual scene to create a coherent percept. Low spatial frequencies convey broad shapes and are preferentially processed by the right hemisphere, while high spatial frequencies convey fine details and are preferentially processed by the left hemisphere (Christman et al., 1991; Mart_í_nez et al., 2001; Peyrin et al., 2004; Piazza & Silver, 2014; Woodhead et al., 2011). Smaller numerosities, characterized by lower spatial frequencies, would engage the right hemisphere and induce a leftward bias. Conversely, larger numerosities, containing higher spatial frequencies, would engage the left hemisphere, guiding attention toward the right. This model provides a plausible biological mechanism for the spatialization of numerosities and predicts the rightward bias induced by smaller physical sizes considering their higher spatial frequencies range with respect to the larger reference stimulus. The absence of a significant leftward bias for the larger stimulus in Experiment 2 may be explained by a conflict between mechanisms driven by spatial frequency and emotional valence, which could cancel the spatial preference. While BAFT predicts that the low spatial frequencies of a larger square should preferentially engage the right hemisphere and induce a leftward bias, the high physical salience of the large red square, which acts as an appetitive stimulus in a rewarded foraging task, may simultaneously trigger a positive emotional response. According to the Emotional-Valence model, large magnitudes associated with a reward recruit the left hemisphere to guide approach behavior toward the right hemispace. These two opposing spatial behaviors, the BAFT-driven leftward bias and the valence-driven rightward approach, may cancel one another, resulting in the observed chance-level performance for larger stimuli.

Our results in day-old chicks, closely mirroring those obtained in 8-to 9-month-old human infants, do not support the prediction that continuous magnitudes follow an innate, congruent small-left/large-right mapping. According to the ATOM (Walsh, 2003) and the “sense of magnitude” theories (Leibovich et al., 2017), a generalized system should ensure that any prothetic dimension is mapped onto space in a similar, congruent manner. However, the presence of a robust spatial-*numerical* association in Experiment 1 and a non-congruent spatial-*quantity* association in Experiment 2 suggests a different developmental priority.

Here, we propose that the mapping of continuous quantity onto space is not an inborn feature but is instead constructed by scaffolding onto a primary, innate capacity to spatialize discrete numerosities. In this view, the “mental number line” acts as the foundational representational framework, and other continuous dimensions only later “borrow” this spatial structure as the organism internalizes environmental correlations between different magnitudes. The fact that even 8/9-month-old human infants, who have already accumulated significant postnatal experience, still lack a congruent spatial-*quantity* association suggests that this integration is a relatively late developmental milestone. To test this hypothesis, future studies could use animals reared under controlled postnatal conditions and test them at different developmental stages to identify when continuous magnitudes begin to align with spatial–numerical mappings. They could also examine whether the strength of spatial–quantity biases covaries with the strength of spatial–numerical biases within individuals. Such studies would clarify whether spatial–quantity associations emerge as a later generalization of spatial–numerical associations rather than as a primitive feature of the cognitive system.

## Acknowledgements

The Authors would like to thank Veronica Pini for her contribution to data collection.

## Author contributions

Conceptualization: A. F., L. R., R. R.; Methodology: L. R., R. R.; Investigation: L. R., R. R; Data analysis: A. F.; Visualization: A.F., Writing-original draft: A. F.; Writing-review and editing: A. F., L. R., R.R.; Funding acquisition: R. R.

## Data availability

Data are publicly available at: https://osf.io/5eu4x/overview?view_only=316d033fb4c54564b9d6e0dc6d120b95

